# The CD36 scavenger receptor Bez regulates lipid redistribution from fat body to oocytes in Drosophila

**DOI:** 10.1101/2023.11.21.568029

**Authors:** Pilar Carrera, Johanna Odenthal, Katharina S. Risse, Yerin Jung, Lars Kuerschner, Margret H. Bülow

## Abstract

Class B scavenger receptors of the CD36 family are important for lipid mobilization from mammalian adipose tissue. This protein family has 3 members in mammals, but 14 in *Drosophila melanogaster*. Lipid distribution in *Drosophila* is mediated by homologs of the LDL receptors, while little is known about the function of scavenger receptors for this process in invertebrates. Here we unravel a role for the so far uncharacterized scavenger receptor Bez in lipid export from *Drosophila* adipocytes. Bez shares the lipid binding residue with CD36 and is expressed at the plasma membrane of the embryonic, larval and adult fat body. Bez loss-of-function lowers the organismal content of storage lipids while they accumulate in the fat body, concomitant with female sterility and degeneration of ovaries. Using an alkyne-labeled fatty acid tracer, we demonstrate that Bez interacts with the apoB homolog Lipophorin at the plasma membrane of adipocytes, thereby enabling lipid transfer. Our study shows how a scavenger receptor interacts with lipoproteins to distribute storage lipids from the fat body to the ovaries and thereby contributes to the metabolic control of development.

## Introduction

Scavenger receptors (SR) are a family of transmembrane proteins that are structurally diverse and have a wide range of ligands. They were first discovered in macrophages as receptors for oxidized Low Density Lipoprotein (oxLDL) (Goldstein et al., 1979; Brown et al., 1980). Further studies showed that SR act as multifunctional receptors implicated in a range of cellular processes. Apart from modified lipoproteins like oxLDL, many other endogenous, mainly polyanionic ligands have been identified, such as long chain fatty acids, oxidized phospholipids and extracellular matrix proteins (Rigotti et al., 1995; Silverstein and Febbraio 2009). SR can be sorted in eight different classes depending on their predicted structure, which is determined by the presence of different conserved motifs (Krieger 1997; Plüddemann et al., 2007). Class B scavenger receptors (SR-B) are characterized by a hairpin structure, formed by two transmembrane domains and one extracellular loop with multiple glycosylation sites. The extracellular CD36-domain includes numerous ligand-binding sites (Canton et al., 2013) and its sequence is highly conserved between the different members of this SR class. Its tertiary structure is defined by disulfide bonds between conserved cysteine residues, which are essential for the recognition of ligands (Gruarin et al., 1997). By contrast, the sequences of the relatively short, intracellular N- and C-terminal tails are not conserved. These tails interact with cytosolic proteins and regulate cellular signalling. SR-B can bind to a wide range of ligands, such as lipoproteins, long chain fatty acids and oxidized phospholipids. In contrast to the other SR classes, they are also able to bind to native lipoproteins such as LDL and HDL (high density lipoprotein) and thereby contribute to the regulation of lipid metabolism. Vertebrate SR-Bs comprise the proteins Cluster of Differentiation 36 (CD36), Scavenger receptor class B member I (SRBI) and Lysosomal integral membrane protein II (LIMP-II). CD36 and SRBI are located at the plasma membrane while LIMP-II is a protein of the lysosomal membrane with a function in the import of glucocerebrosidase (Heybrock et al., 2019).

CD36 is a multifunctional protein that regulates the uptake and export of fatty acids in gut cells, adipocytes and muscle cells (Daquinag et al., 2021). It is also expressed in taste bud cells where it acts as a lipid sensor for the detection of dietary lipids (Laugerette, 2005). Fatty acid transfer from adipocytes to ovarian tumours has been shown to depend on the CD36-dependent lipid uptake by adipocytes (Ladanyi et al., 2018). CD36 is also required for ovarian angiogenesis and folliculogenesis: CD36-deficient mice display hypervascularized ovaries and have less offspring than controls (Osz et al., 2014). However, this function has been linked to binding of thrombospondin-1, an angiogenesis regulator, and not to the lipid binding property of CD36.

In the invertebrate *Drosophila melanogaster*, 14 SR-B homologues have been identified (Nichols & Vogt, 2008). The high number of SR genes suggests a diversification of functions in insects. Many of these genes remain uncharacterized until now, but some have been functionally analysed. Croquemort (Crq) is expressed in macrophages and takes part in the phagocytosis of apoptotic cells during embryogenesis and the morphogenesis of the central nervous system (Guillou et al., 2016). Debris buster (Dsb) is a protein of the lysosomal membrane that is required for trafficking of extracellular matrix components important for airway physiology and for clearing of degenerating dendrites (Han et al., 2014; Wingen et al., 2017). Neither inactivation nor afterpotential-D (NinaD) and Scavenger receptor acting in neural tissue and majority of rhodopsin absent (Santa-Maria) were identified as transporters for carotenoids, precursors for the synthesis of vitamin A, which itself is modified and then embedded in the optical pigment Rhodopsin. NinaD is expressed in the gut where it mediates the uptake of carotenoids, which are required for visual chromophore synthesis. Santa-Maria is expressed in specific neurons and glia cells and locates to the plasma membrane, where it mediates the uptake of β-carotene that afterwards is converted to vitamin A (Dewett et al., 2021). Sensory neuron membrane protein 1 (SNMP1) is expressed in olfactory neurons and in steroidogenic organs such as the ovaries, testes and the prothoracic gland (PG). Silencing of SNMP1 via RNAi expression in the PG and in the follicle cells of the ovary results in a reduced content of lipid droplets in these tissues (Talamillo et al., 2013).

Lipid shuttling in both human blood and Drosophila hemolymph is mediated by lipoproteins. In Drosophila, apoLpp (Lipophorin) and apoLTP (Lipid transfer protein) belong to the apoB family of lipoproteins. The product of the *apoLpp* gene (CG11064) gives rise to apoLI and apoLII that assemble the Lipophorin lipoprotein. Human apoB is the major lipoprotein in LDL particles. Lpp binds to LTP in the gut and is loaded with dietary lipids. These are distributed to the fat body for storage and to other organs (Palm et al., 2012). The Drosophila fat body stores lipids in lipid droplets from where they are mobilized, e.g. upon starvation, to nurse other tissues. Fasting induces a morphological change of fat body lipid droplets as they become larger and fewer when lipids are mobilized (Ugrankar et al., 2019).

Nutrition is closely linked to reproduction. Malnutrition in women heavily impacts reproduction, with both obesity and undernutrition affecting fertility (Silvestris et al., 2019). Also in *Drosophila*, this link exists and Drosophila females store high amounts of triglycerides (TG), a neutral lipid known as fat to support oocyte maturation. Uptake of neutral lipids takes place between stage 8 and 10 of oocyte maturation: at stage 10, oocytes massively accumulate neutral lipids in both nurse cells and follicle cells. Lipoproteins mediate the transfer of lipids from the hemolymph to ovaries (Sieber & Spradling, 2015). In ovaries, lipophorin receptors bind LTP, which is required for neutral lipid accumulation (Rodríguez-Vázquez et al., 2015). The metabolic signalling that allows the transfer of storage lipids from adipocytes to ovaries is still incompletely understood. Here we identify CG3829 as one of the few of the 14 Drosophila SR-B that contain a conserved lipid binding residue homologous to CD36. We show its role for the export of dietary lipids from the fat body to the circulation. We named the protein Bez (**B**etween **E**mp and **Z**ip, pronounce “Beth”) and found that in Bez mutants, ovaries degenerate due to missing lipid transfer from the fat body. Bez mutant adipocytes accumulate lipids and are unable to mobilize TG, and Bez-mediated lipid export from the fat body is required to allow lipid accumulation during oocyte maturation. We demonstrate that Bez transfers labeled lipids to the lipoprotein apoLpp, thereby distributing storage lipids to ovaries. We thus establish a role for this SR-B in the metabolic regulation of reproduction.

## Results

### The class B scavenger receptor Bez is expressed at the plasma membrane of adipocytes

The *Drosophila* class B scavenger receptor (SR-B) Bez (CG3829) is one of 14 CD36-like protein family members. We performed an Alphafold analysis of its predicted protein structure (Figure 1A). Similar to human CD36, CG3829 contains two transmembrane domains and a large, extracellular barrel-like putative lipid binding domain with high sequence similarity to CD36 (Supplemental Figure S1). Of note, Bez is the only fly SR-B that contains two conserved lysine residues of the hydrophobic pocket that in mammalian CD36 has been linked to lipid uptake (Kuda et al., 2013). The SR-B Peste (CG7228) and Santa-maria (CG12789) contain one of these lysine residues, which are absent in the other *Drosophila* SR-B. In addition, Bez contains two disorganized amino acid stretches at both termini. We analyzed the expression pattern of the Bez protein in embryos and detected expression at the apical membrane of the embryonic gut (Supplemental Figure S2A) and at the yolk sac (Supplemental Figure S2B). Strong expression was also seen in the fat body (Figure 1B-D). In the embryonic fat body (Figure 1B) marked by the lipid droplet surface protein Perilipin-1 (Plin1, Beller et al., 2010) Bez was found primarily at the plasma membrane of the adipocytes (Figure 1C). To confirm this localization we generated bez-RNAi clones in the fat body. Wildtype cells (GFP-negative) show high Bez expression at the plasma membrane (illustrated by colocalization with the plasma membrane protein alpha-Spectrin), while bez-RNAi clones (marked by the GFP signal) lacked Bez staining (Figure 1C, asterisks). Bez also localizes to the plasma membrane of the adult fat body adipocytes, marked by Plin1 (Figure 1D). Taken together, the Bez expression pattern suggests a conserved function in lipid uptake similar to that of CD36.

**Figure 1:**
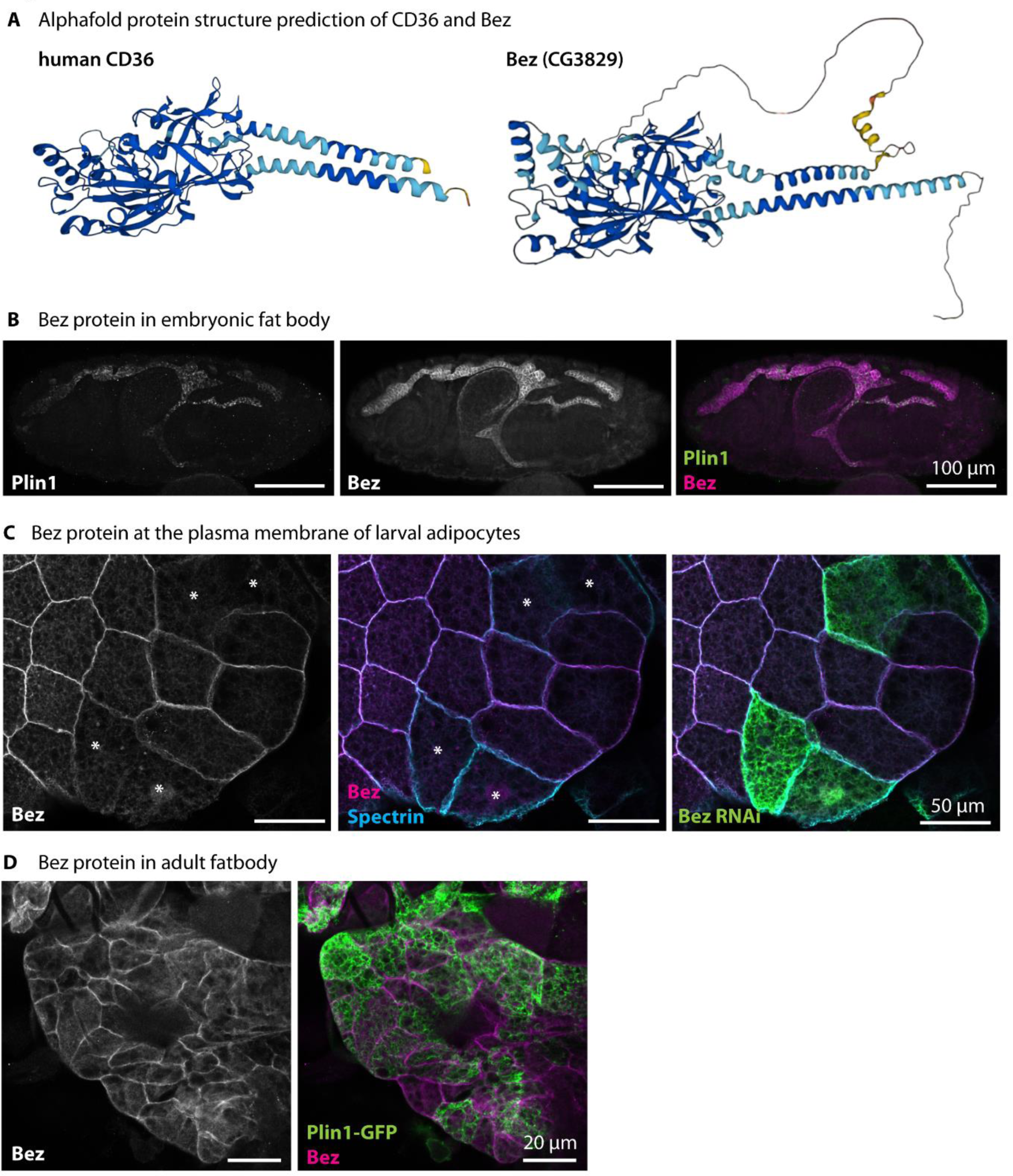
The class B scavenger receptor Bez is expressed at the plasma membrane of adipocytes. A) Protein structure prediction by Alphafold of human CD36 (Platelet glycoprotein 4) and *Drosophila melanogaster* CG3829 (Bez) shows the N- and C-terminal transmembrane domains and the extracellular, barrel-formed lipid transport domain of CD36. The two transmembrane domains and the extracellular, putative lipid transport domain are conserved in *D. melanogaster*. In addition, Bez has two disorganized amino acid stretches at the N- and C-terminus. B) The Bez protein is expressed in the embryonic fat body: Confocal images of embryos show co-expression of Bez and the lipid droplet protein Plin1. Representative images from 5 independent replicates. Scale bars represent 100 µm. C) The Bez protein is expressed at the plasma membrane of larval fat body cells. Confocal images of 3^rd^ instar larval fat body shows colocalization of Bez with the membrane marker alpha-Spectrin. Clones expressing bez-RNAi are marked with GFP and show no Bez staining in the plasma membrane. Representative images from 5 independent replicates. Scale bars represent 50 µm. D) Bez is expressed at the plasma membrane of adult fat body. Plin1-GFP was expressed in adipocytes using cg-Gal4. Representative images from 5 independent replicates. Scale bars represent 20 µm.

### Bez in the fat body is required for developmental timing

To investigate the molecular function of Bez, we used a mutant where the P-element P(EP)CG3829^G8378^ was inserted into the open reading frame (mutant allele *bez^EP^*). By mobilizing this EP element we generated a second allele (*bez^jo2^*) featuring a stop codon in the sequence encoding for the first transmembrane domain at the position of amino acid 89 (Figure 2A). The resulting Bez mutants (*bez^EP/jo2^*, bez-/-) do not express the Bez protein in embryos (Supplemental Figure S3). Bez homozygous mutants are semi-viable, but females are sterile, and larvae show a strong developmental delay. 50% of heterozygous controls pupariate after ∼4,5 days at 25°C, while 50% of homozygous Bez mutants pupariate at almost 6 days after egg laying (AEL). Similarly, adult flies hatch ∼1,5 days later than the heterozygous controls (Figure 2B). Expression of the UAS-Bez construct in the fat body (*cg*-Gal4) reversed the developmental delay of Bez mutants (Figure 2B, “rescue”). Next, we expressed RNAi against Bez in different tissues. Expression of Bez RNAi in the midgut (*mex*-Gal4) or in hemocytes (*hml*-Gal4) did not phenocopy the developmental delay of Bez homozygous mutants. By contrast, expression in the fat body (*cg*-Gal4) fully phenocopied the developmental delay (Figure 2C). This shows that Bez function in the fat body is essential for normal development.

**Figure 2:**
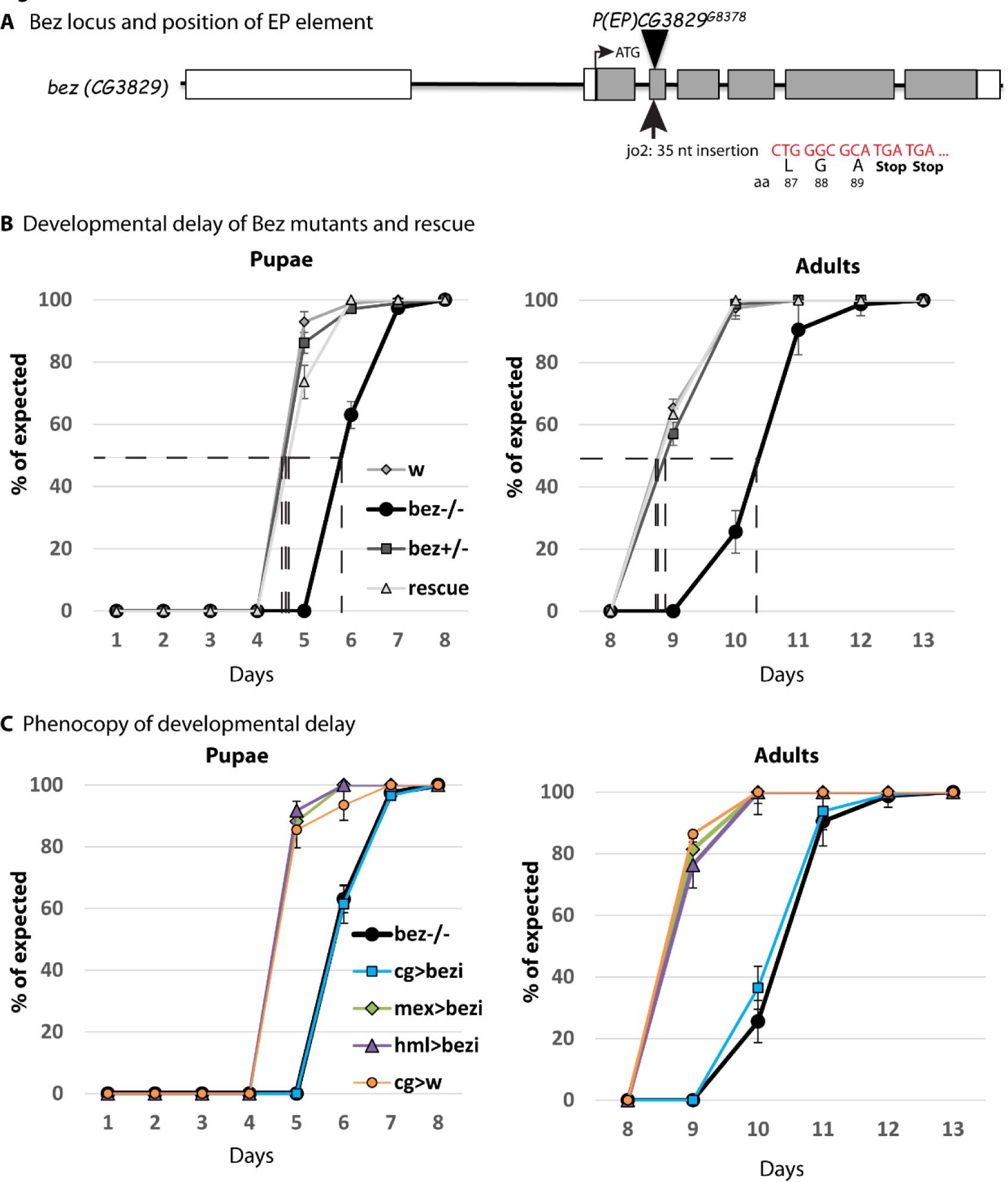
Bez is required in the fat body for developmental timing. A) Representation of the *bez* locus and the position of the EP element that mutate the *bez* locus and was used to create an additional jump-out (jo2) allele. Nucleotide insertion contained in the jo2 allele is shown in red. The 35 nt insertion yields two stop codons after amino acid 89. B) Bez mutant development is delayed by >1 day. 50% of heterozygous controls pupariate after 4.5 days at 25°C while homozygous mutants pupariate after 5.75 days. Adult hatching is similarly delayed, with 50% of heterozygous controls hatching after 9 days and homozygous mutants after 10.3 days. Developmental delay is rescued by expression of UAS-Bez in the fat body (*cg*-Gal4). Error bars represent SEM. n=5 in groups of 20 individuals. C) The delay in pupal and adult development is phenocopied by knocking down Bez in the fat body (cg-Gal4, blue line, control in orange) by RNAi, but not in the midgut (mex-Gal4, green line) or in hemocytes (hml-Gal4, purple line). Error bars represent SEM. n=5 in groups of 20 individuals.

### Bez function in adipocytes is essential for female fertility

To analyze the cause for female sterility of Bez mutants, we investigated Bez mutant fecundity, egg morphology, and ovary morphology. We found that Bez mutants lay fewer and smaller eggs and that these eggs collapse, with no larvae hatching from them (Figure 3A). We dissected ovaries of control females and Bez mutant females and found that also ovaries are small and degenerated (Figure 3B). We then asked whether Bez was expressed in ovaries and performed quantitative real-time PCR of RNA isolated from whole flies and from isolated ovaries (Figure 3C). We found that Bez is not expressed in ovaries, suggesting that Bez functions remotely from the fat body to sustain ovary maturation. We analyzed oogenesis by immunofluorescent staining of dissected ovaries and found that oocyte maturation during oogenesis is blocked in Bez mutants: while control ovarioles contain oocytes of all stages including mature eggs, Bez mutant ovarioles contain no mature eggs and no oocytes older than stage 10. Localization of Orb, an RNA-binding protein that establishes polarity during oogenesis (Lantz et al., 1994), was broader and less defined in the degenerated egg chambers of Bez mutants (Figure 3D). Degenerated oocytes were apoptotic and showed fragmented DNA, as shown by the fragmented DAPI staining. Next, we probed for the tissue-specific phenocopy of Bez mutant fecundity. We found that expression of Bez RNAi in the fat body (*FB*-Gal4 or *cg*-Gal4) as well as ubiquitous expression (*c355*-Gal4) is sufficient to block fecundity (Figure 3E). Expression of Bez RNAi in the fat body (*FB*-Gal4 or *cg*-Gal4) was also sufficient to block oocyte maturation and to induce oocyte apoptosis similar to Bez mutants (Figure 3F, Supplemental Figure S4A,B). Our results show that Bez function in adipocytes is essential for oocyte maturation and thereby for female fecundity.

**Figure 3:**
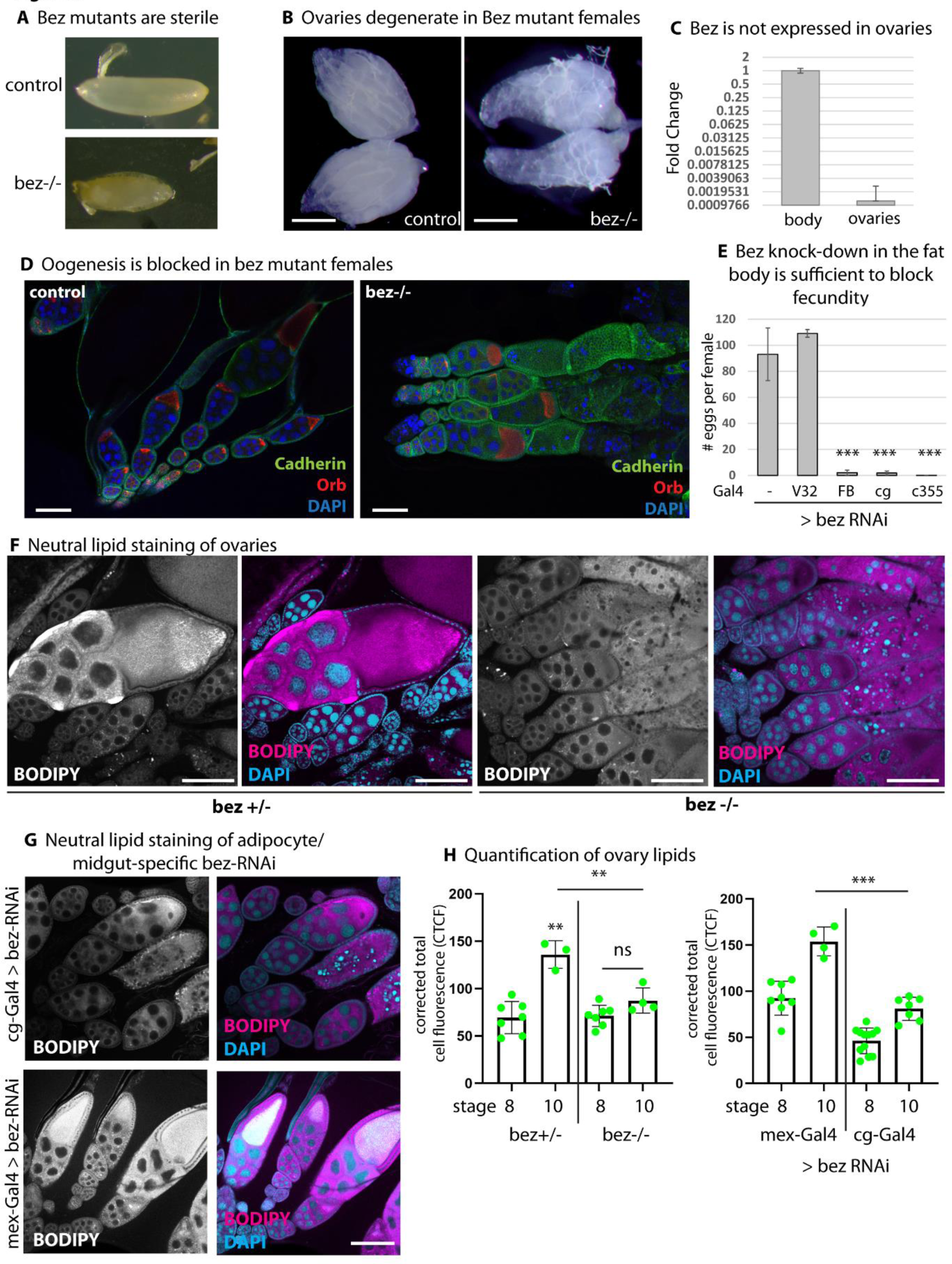
Bez function in adipocytes is essential for female fertility. A) Bez mutant females are sterile and lay unviable, collapsed eggs. Representative images from 10 independent replicates. B) Bez mutant female ovaries degenerate. Representative images from 10 independent replicates. Scale bars represent 250 µm. C) The *bez* transcript is not expressed in ovaries. Quantitative real-time PCR from isolated ovaries compared to whole females. Scale bars represent standard deviation. Mean of 8 experiments. D) Oogenesis is blocked in Bez mutants. Confocal images show wildtype and Bez mutant ovaries stained with anti-E-Cadherin, anti-Orb and DAPI. Bez mutant oocytes do not mature into eggs. Representative images from 10 independent replicates. Scale bars represent 100 µm. E) Knock-down of Bez by RNAi in the fat body (*FB*-Gal4, *cg*-Gal4 and *c355*-Gal4) is sufficient to block fecundity, while knock-down of Bez in germ line (*V32*-Gal4) cells has no effect. n=5 in groups of 20 individuals. Asterisks represent *** p > 0.001. F) Bez mutant ovaries do not store lipids. Neutral lipid staining (BODIPY) shows lipid accumulation in stage ≥ 10 ovaries in heterozygous controls, but not in homozygous Bez mutants. G) Bez RNAi knock-down in the fat body (*cg*-Gal4) is sufficient to reduce the lipid content of ovaries. Bez RNAi knock-down in the midgut (*mex*-Gal4) does not block lipid accumulation on ovaries and does not induce oocyte degeneration. Representative images from 5 independent replicates. Scale bars represent 100 µm. H) Quantification of fluorescence intensity of BODIPY from F and G. Asterisks represent *** p > 0.001, ** p > 0.01, ns: not significant.

### Bez mutants are starvation-sensitive and do not mobilize triglyceride upon starvation

Oocytes display a shift in their lipid content between stage 8 and 10. Stage 10 oocytes store high amounts of neutral lipids (Sieber & Spradling, 2015). To investigate if defects in lipid storage cause ovary degeneration in Bez mutants, we stained neutral lipids with BODIPY. We confirmed that stage 10 oocytes massively take up storage lipids in heterozygous controls (Figure 3F, bez+/-). Degenerated oocytes of homozygous Bez mutants did not contain increased amounts of neutral lipids (Figure 3F, bez-/-). We quantified the fluorescence intensity of BODIPY in egg chambers between stage 8 and 10, which showed that Bez homozygous mutant egg chambers do not increase their lipid content (Figure 3H). This missing lipid uptake was phenocopied by expression of Bez RNAi in the fat body (*cg*-Gal4, Figure 3G, H). By contrast, the stage 8 – stage 10 lipid transition was independent from midgut expression of Bez: midgut depletion of Bez allowed for lipid accumulation, and did not trigger egg chamber degeneration (*mex*-Gal4, Figure 3G, H). Next, we used a colorimetric assay to measure triglycerides (TG) of whole female flies. Fat body-specific depletion of Bez by RNAi led to significantly reduced total TG content compared to controls (Figure 4A).

**Figure 4:**
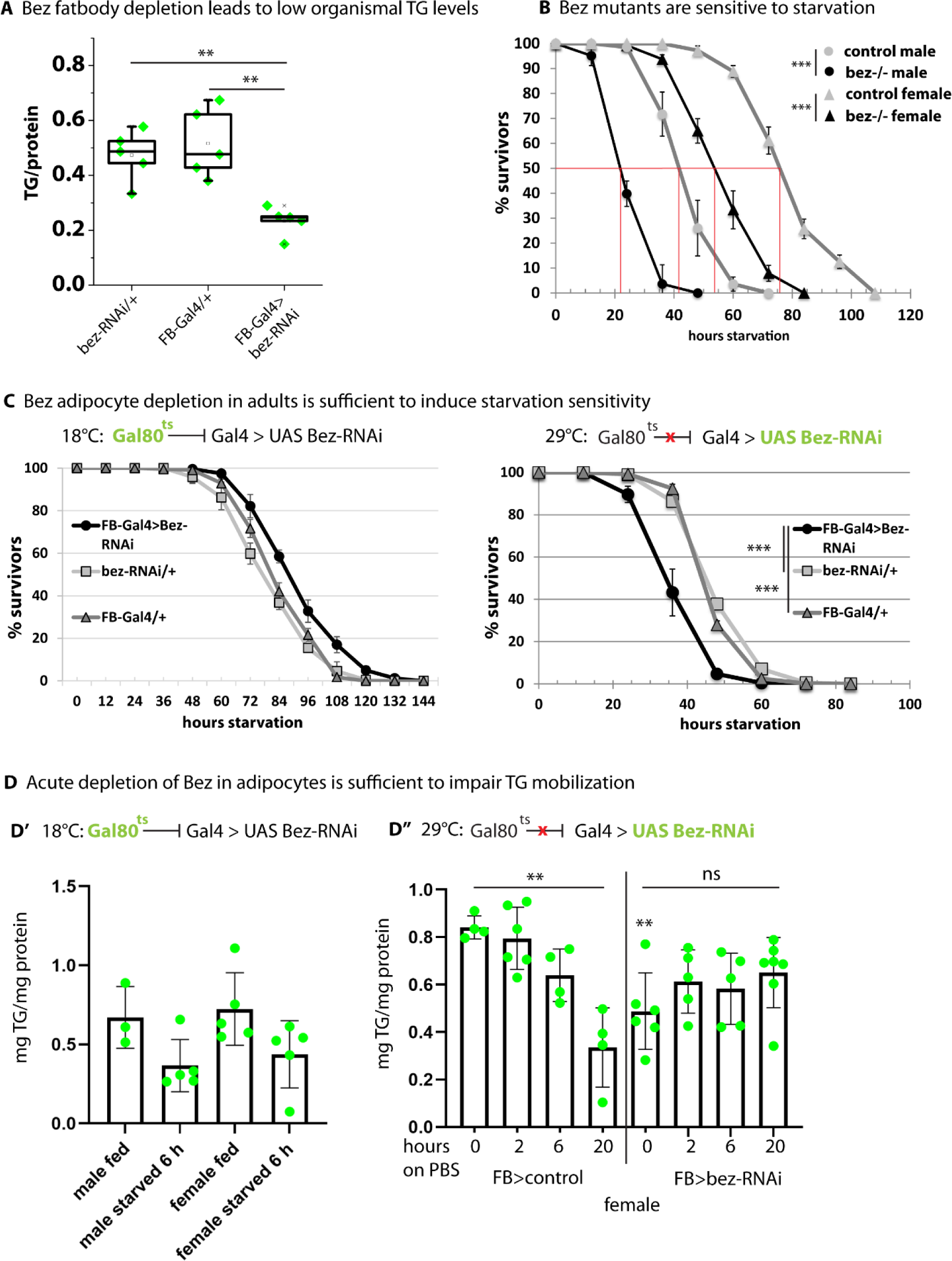
Bez mutants are starvation-sensitive and fail to mobilize lipids. A) RNAi knock-down of Bez in the fat body (*FB*-Gal4) reduces the levels of the storage lipid triacylglycerol as determined by a colorimetric assay. n=5 in groups of 8-10 individuals. Asterisks represent ** p > 0.01. B) Bez mutants are sensitive to starvation. 50 % of wildtype female flies are dead after ∼75 hours under food deprivation while 50 % of Bez mutants are dead after ∼50 h. Male flies have lower starvation resistance: 50 % wildtypes are dead after ∼40 h and ∼20 h in Bez mutants. Flies were kept at 25 °C. n=5 in groups of 20 individuals. Asterisks represent *** p > 0.001. C) Adult onset of Bez fat body depletion is sufficient to induce starvation sensitivity in females, suggesting that it is due to lipid mobilization defects rather than storage of lipids. bez-RNAi was expressed in the fat body (*FB*-Gal4). The thermosensitive Gal80 inhibitor was blocked in adults at 29°C, leading to Bez depletion and starvation sensitivity. n=5 in groups of 20 individuals. Asterisks represent *** p > 0.001. D) Triacylglycerol (TG) as determined by a colorimetric assay from whole flies expressing bez RNAi in the fat body (*FB*-Gal4) under the control of the inducible Gal4/Gal80 system. At 18 °C, Gal4 is inactive and both male and female flies can mobilize TG upon starvation (D’). At 29 °C, Gal4 is active and drives the expression of bez-RNAi (D’’, right side) in female flies. In controls, FB Gal4/Gal80^ts^ was crossed to wildtype (D’’, left side). Depletion of Bez from the fat body blocks TG reduction upon starvation. n ≥ 3 in groups of 8 individuals. Asterisks represent ** p > 0.01, ns: not significant.

We hypothesized that Bez is required for lipid mobilization from adipocytes, and that under Bez loss-of-function, impaired lipid mobilization leads to insufficient lipid supply of oocytes. Lipid mobilization as well as oocyte supply with lipids is strongly dependent on nutrient availability.

We thus asked if Bez mutants are unable to mobilize lipids upon starvation. Control female flies can survive up to four days without food, with 50 % of flies surviving after 76 hours of starvation. Male flies are less starvation-resistant with 50 % of flies surviving after 40 hours of starvation (Figure 4B). Both female and male Bez mutants are significantly less starvation-resistant than controls. To investigate if acute depletion of Bez from the fat body is sufficient to trigger starvation sensitivity, we expressed Bez RNAi under the control of the temperature-sensitive Gal80 system in females. At 18°C, Gal80 expression suppresses Gal4 and thus the expression of bez-RNAi. At 29°C, Gal80 no longer represses Gal4, allowing for the expression of bez-RNAi in the fat body. The resulting acute depletion of Bez in adipocytes induced starvation sensitivity compared to controls (Figure 4C). Next, we repeated the experiment and measured TG levels in whole male and female flies. At 18°C, the TG content is reduced in response to nutrient restriction in both male and female flies (Figure 4D’). At 29°C, Bez depletion in adipocytes reduced the organismal TG content in both male and female flies (Figure 4D’’) and control flies reduced their TG content in response to nutrient restriction. TG levels in male (Supplemental Figure S5A) and female (Figure 4D’’) flies with Bez depletion in adipocytes remained at similar levels under starvation. This shows that acute and adipocyte-restricted depletion of Bez impairs lipid mobilization upon starvation.

### Bez loss-of-function alters lipid droplet size and lipid uptake into adipocytes

We hypothesized that impaired lipid mobilization in Bez mutants would affect lipid droplet morphology. We used larval fat body tissue of control animals and Bez mutants and stained lipid droplets with BODIPY. The lipid droplet size was semi-quantified as the BODIPY-stained area (Figure 5A, B). Bez mutant fat body tissue contained significantly enlarged lipid droplets, while their number per cell remained unchanged. Of note, this matched the lipid droplets phenotype in adipocytes of loss-of-function mutants of Lipophorin (Lpp), a Drosophila lipoprotein homologous to ApoB (Palm et al., 2012). We repeated the experiment in Bez mutant fat body tissue with Bez reconstituted in distinct clones allowing for direct comparison of lipid droplet morphology within the same tissue (Supplemental Figure 6A, B). Lipid droplet staining confirmed that these organelles are large in Bez mutant adipocytes, while in adjacent rescue clones the droplets are small and more numerous. Similarly, bez-RNAi leads to larger lipid droplets (Supplemental Figure S6C). Importantly, overexpression of Bez using the UAS-Bez rescue construct did not alter lipid droplet morphology in a wildtype background (Supplemental Figure S6D).

**Figure 5:**
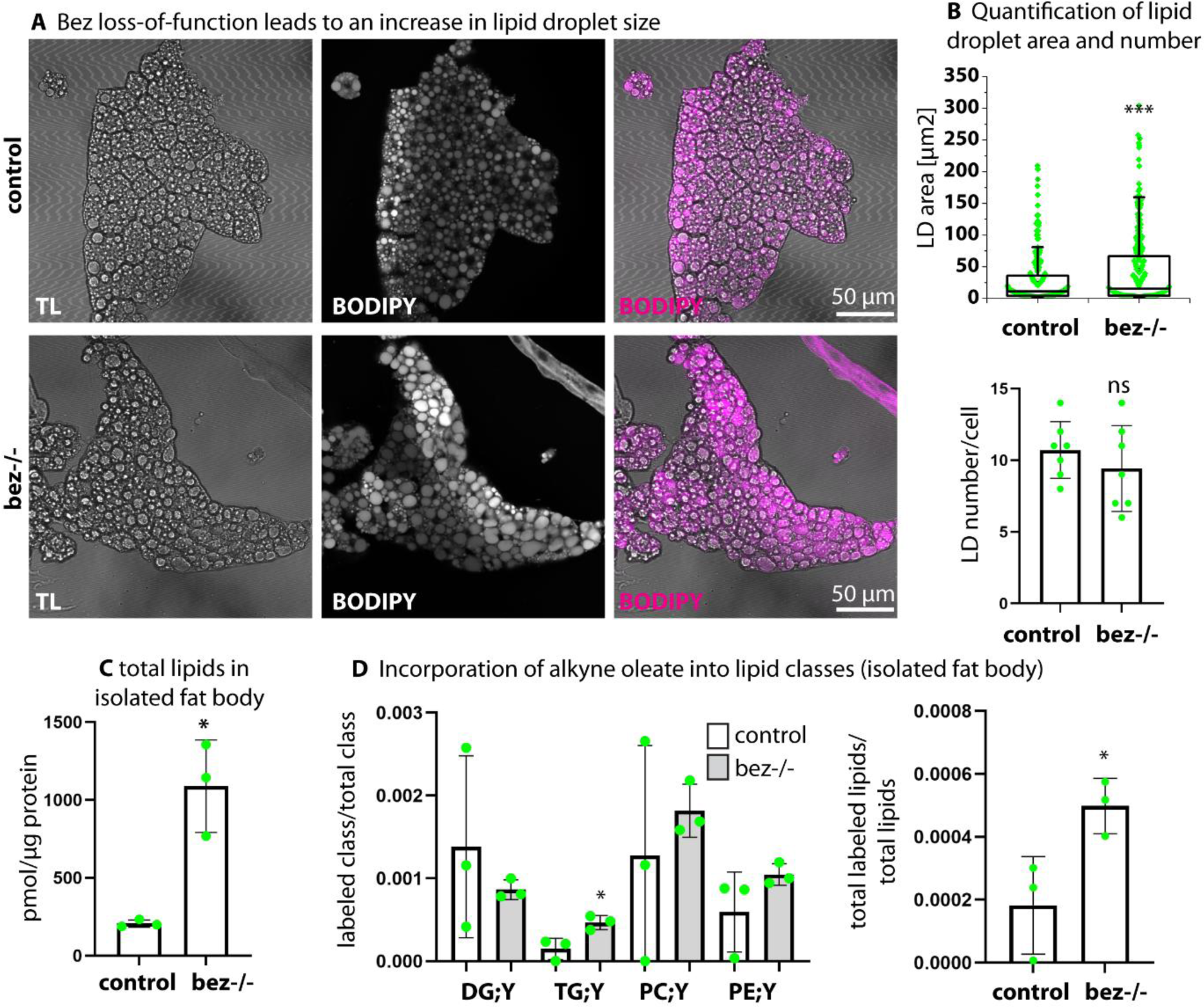
Bez loss-of-function affects lipid droplet size and lipid uptake. A) Confocal images show control (w-) and Bez mutant adipocytes in fat body tissue. TL: transmission light. Lipid droplets are stained with BODIPY. Bez mutant cells show increased lipid droplet size. Representative images from 10 independent replicates. B) Quantification of lipid droplet size (area) and number per cell. Asterisks represent *** p > 0.001, ns: not significant. C) Lipid content of 5 isolated fat bodies determined by mass spectrometry and normalized to protein content. n = 3. Asterisk represents * p > 0.05. D) Incorporation of alkyne oleate (FA19:1;Y) into lipid classes as determined by click reaction followed by mass spectrometry. Shown are incorporation into diacylglycerol (DG;Y), Triacylglycerol (TG;Y), phosphatidylglycerol (PC;Y), phosphatidylethanolamine (PC;Y) and total lipids. n = 3. Asterisk represents * p > 0.05.

Next, we asked if the difference in lipid droplet morphology in Bez mutant fat body tissue corresponds to differences in lipid content. We determined the total lipid content of isolated fat body tissue of control and Bez mutant larvae by mass spectrometry (Figure 5C). Bez mutants accumulate lipids of both types, the neutral lipids (DG, TG) and phospholipids (PE, PC, Supplemental Figure 6E). To investigate how Bez affects uptake and incorporation of fatty acids into the lipidome of the fat body we incubated isolated tissue of control flies and Bez mutants with alkyne-labeled oleic acid (alkyne oleate, FA 19:1;Y) a traceable analog of the abundant natural oleic acid (Kuerschner & Thiele, 2022). Upon click-reaction a mass spectrometry lipidome analysis was performed (Wunderling et al., 2021). Bez mutants showed significantly increased total uptake and incorporation of alkyne oleate into the TG pool generating labeled TG;Y (Figure 5D). Levels of labeled phospholipids (PE;Y and PC;Y) were also elevated in the mutant tissue, but our measurements did not reach significance. Taken together, this data shows that Bez mutant adipocytes enhance lipid uptake and storage, but are unable to mobilize these lipids. This suggests that lipid export from adipocytes to target tissues (such as oocytes) is impaired in Bez mutants.

### Bez interacts with the apolipoprotein Lipophorin to distribute storage lipids

How does Bez affect the lipid content of oocytes without being present in ovaries? Lipids are transported between organs by lipoproteins in both *Drosophila* and humans. A homolog of ApoB is Rfabg/apoLpp (Lipophorin). We used a fly line that expresses Lpp fused to GFP under the control of its endogenous promoter. We first analyzed if Bez colocalized with Lpp at the adipocyte plasma membrane. Using airyscan super-resolution microscopy we found that both Bez and Lpp are organized in distinct punctate domains at the adipocyte plasma membrane, several of which colocalize (Figure 6A, Supplemental Figure 7A). Next, we asked if Bez has an impact on the lipid composition of the lipoprotein particles. We isolated particles from Rfabg reporter animals of wildtype or Bez mutant background and analyzed their lipid content by mass spectrometry. Although similar in protein content (Figure 6B), particles from Bez mutants show a strongly reduced lipid content, just missing statistical significance (Figure 6C). We then assessed the distribution of Lpp in the gut and fat body. The apical membrane of the gastric caeca was marked by Lpp-GFP in control animals, while in *Bez* mutants this membrane showed an even more intense signal (Figure 6D). In the fat body, Lpp-GFP is found in a regular pattern at the plasma membrane of adipocytes, similar to the Bez protein (Figure 6E). In the absence of Bez, this regular pattern is lost and Lpp-GFP presents in aggregates at the plasma membrane.

**Figure 6:**
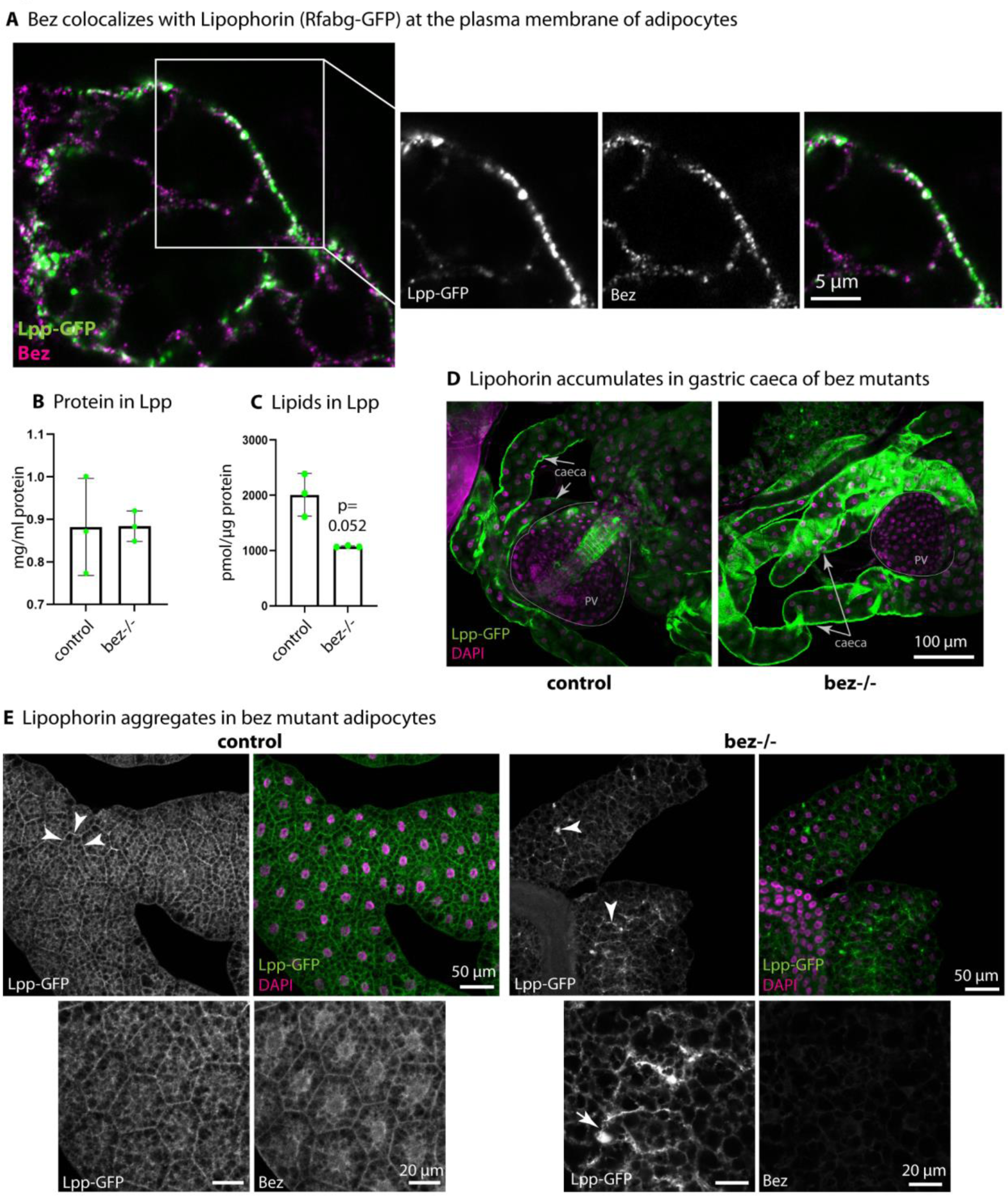
Bez interacts with the apolipoprotein apoLpp. A) Larval fat body cells from Rfabg (apoLpp)-GFP expressing animals (under the control of an endogenous promoter). Airyscan confocal images with the endogenous Bez scanned in the AiryScan mode show Lpp-GFP and Bez colocalization at the plasma membrane of adipocytes. Images show single confocal section. Representative images from 5 independent replicates. Scale bar as indicated. B) Protein content of isolated Lpp fraction of control and Bez mutant larvae. n = 3. C) Lipid content of isolated Lpp fraction as determined by mass spectrometry and normalized to protein. n = 3. D) Confocal images of larval anterior midgut showing gastric caeca and proventriculus (PV) of Lpp-GFP and Lpp-GFP; Bez-/- larvae. Representative images from 5 independent replicates. Scale bar as indicated. E) Confocal images of larval fat body of Lpp-GFP and Lpp-GFP; Bez-/-. Lpp localizes in a regular pattern at the plasma membrane and colocalizes with Bez in controls. In Bez mutants, Lpp forms aggregates and no longer marks the plasma membrane. Representative images from 5 independent replicates. Scale bars as indicated.

Lipophorins were found to have a distinct fatty acid profile: in DG, medium-chain fatty acids (MCFA) with an average chain length of 12-14 carbons are abundant, while in phospholipids, fatty acids with an average chain length of 16-18 carbons are prevalent (Palm et al., 2012). Loading of MCFA-DG to Lpp depends on transfer by lipid transfer particle (LTP) in the gut (Palm et al., 2012). We deepened our mass spectrometry analysis of the Lpp lipid content by evaluating the fatty acid distribution within the pools of DG and phospholipids (Supplemental Figure S7B). We confirmed the presence of MCFA-DG and phospholipids with an average chain length of 16-18 carbons in isolated Lpp by mass spectrometry. Strikingly, in Bez mutants a subpopulation of DG species containing medium-chain fatty acids with chains of 12-14 carbons was found reduced and the total DG pool shifted towards longer fatty acid chains (Supplemental Figure S7B). The fatty acid profile of phospholipids was also slightly shifted, but fatty acids with an average chain length of 16-18 carbons remain the most abundant species in phospholipids of Bez mutant Lpp. Together, this suggests that Bez might be required for LTP/Lpp-mediated lipid export from the gut and for correct interaction of Lpp with the adipocyte plasma membrane. However, knock-down of Bez in the gut did not trigger a developmental delay or block oocyte maturation (Figure 2C, Figure 3G, H). We thus hypothesize that interaction of Bez and Lpp in the fat body determines lipid delivery to oocytes and is limiting for oocyte maturation.

### Bez mediates lipid transfer from adipocytes to Lipophorin

As Bez localizes to the plasma membrane of adipocytes it becomes exposed to hemolymph and lipoprotein particles. To corroborate the interaction of Lpp from the hemolymph with the membrane receptor Bez, we isolated hemolymph from Lpp-GFP-expressing flies for incubation with Bez mutant fat body tissue that contains Bez rescue clones (Figure 7A). In fat bodies incubated with hemolymph from wildtype control flies, only background staining was visible in the GFP channel (Figure 7A, upper panels). Alike, Bez mutant cells incubated with hemolymph from Lpp-GFP flies show only minor signal for GFP (Figure 7A, lower panels). By contrast, the rescue clones show a strong staining for apoLpp-GFP demonstrating a Bez-dependent recruitment of Lpp to the plasma membrane of adipocytes.

**Figure 7:**
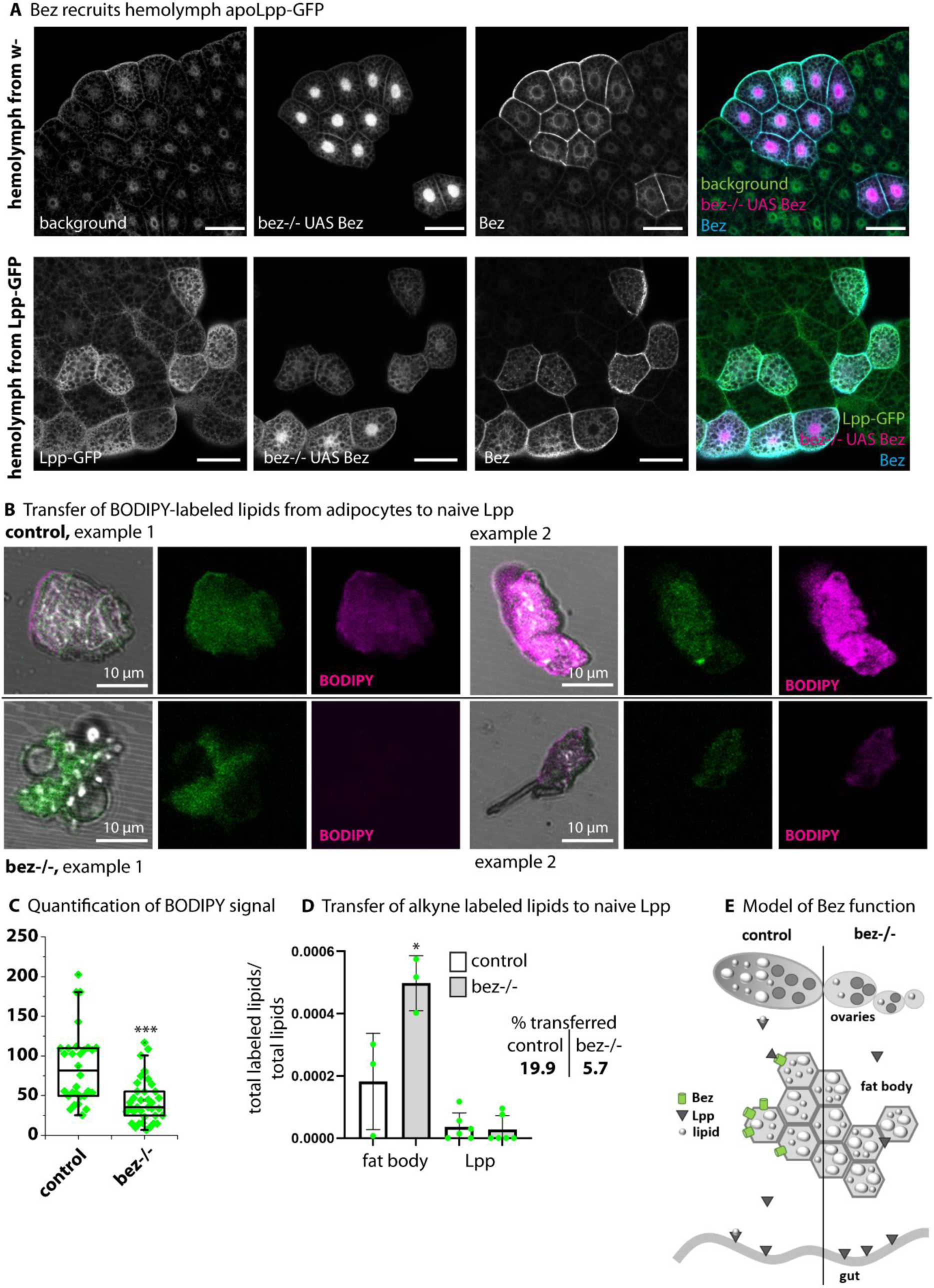
Bez is required for lipid transfer from adipocytes to Lpp. A) Bez can recruit the lipoprotein Rfabg (apoLpp, CG11064) to adipocytes. The upper panel shows the control that was incubated with hemolymph from w-controls. Bez rescue clones are labeled with RFP, RFP-negative cells are bez-/-. The lower panel shows bez-/- fat body tissue that was incubated with hemolymph from Lpp-GFP-expressing flies. Bez rescue clones are marked with RFP. Bez-/- show only background staining for Lpp-GFP, while rescue clones strongly bind Lpp. Representative images from 5 independent replicates. Scale bars represent 50 µm. B) Confocal images of Lpp-GFP aggregates. Lpp from the Rfabg line that expresses Lpp-GFP under its endogenous promoter were incubated with BODIPY-stained fat body from controls and Bez mutants. BODIPY is readily transferred from control adipocytes to naïve Lpp (upper panel, two examples are shown). Transfer of BODIPY-labeled lipids is absent (example 1) or reduced (example 2) from Bez mutant adipocytes. Representative images from 5 independent replicates. Scale bars as indicated. C) Quantification of the BODIPY fluorescence intensity in Lpp aggregates from B. Asterisks represent *** p > 0.001. D) Transfer of clicked alkyne oleate as determined by mass spectrometry. Lpp from the Rfabg line that expresses Lpp-GFP under its endogenous promoter were incubated with alkyne oleate-loaded fat body from controls and Bez mutants. Bez mutant adipocytes incorporate more alkyne oleate than control adipocytes, but hardly transfer the traceable analog to Lpp. E) Model of Bez function: in the presence of Bez, lipids can be transferred from adipocytes to Lpp and distributed to ovaries. Lipids accumulate in the fat body and ovaries cannot mature in Bez mutants.

We asked if the lipid transfer to Lpp by adipocytes depends on Bez. To address this question, we used BODIPY to stain neutral lipids in control and Bez mutant fat bodies. Isolated Lpp-GFP particles not containing any lipid dye were incubated for 60 min with the stained fat bodies. Upon removal of the tissue the Lpp-GFP particles were analyzed by microscopy (Figure 7B). Lpp-GFP is visible as small, GFP-positive aggregates. Lpp that were incubated with control fat body readily incorporated the BODIPY dye. By contrast, Lpp exposed to fat body of Bez mutants showed a significantly reduced BODIPY signal (Figure 7B, C). In a parallel set of experiments omitting the BODIPY stain we used the alkyne oleic acid instead (Figure 7D). Here the different fat bodies were pre-incubated with tracer yielding labeled lipid metabolites. After washing, the tissue was incubated for 60 min with naïve Lpp before separation of tissue and Lpp. The particles were washed and analyzed by mass spectrometry for their content of labeled lipids. As shown before (Figure 5D), Bez mutant adipocytes readily incorporate alkyne oleate into their lipidome (Figure 7D, left). However, despite their elevated content of labeled lipids, Bez mutant adipocytes struggle to transfer labeled lipids to Lpp (Figure 7D, right). Accordingly, naïve Lpp incubated with Bez mutant tissue acquire relatively lower amounts of labeled lipids: only ∼6% were transferred, compared to ∼20% for controls. Taken together, these experiments show that Bez is required for productive lipid transfer from fat body to Lpp.

In sum, we suggest that Bez is required for lipid export from adipocytes, and that by interaction with lipoproteins it regulates the distribution of lipids to ovaries, where these lipids are required for oocyte maturation (Figure 7E). We thus demonstrate a role for a previously uncharacterized scavenger receptor in the remote metabolic regulation of ovary development.

## Discussion

The CD36 scavenger receptor plays a crucial role in fatty acid transport. CD36 is required to incorporate dietary lipids into chylomicrons, which allows the transfer of fatty acids from the gut into the circulation. Fatty acids transported by LDL are taken up into adipose tissue by LDL receptors, and mobilization of lipids from adipocytes again requires CD36 (Daquinag et al., 2021). However, whether CD36 interacts with lipoproteins in this process is unresolved yet. CD36 is also required for adipogenesis, and patients with a CD36 deficiency show impaired chylomicron formation, reduced lipid utilization and storage, and increased lipolysis (Zhao et al., 2018). Here we describe a novel role for Bez, a largely uncharacterized CD36-like lipid scavenger receptor in lipid export from adipocytes of *Drosophila melanogaster*. We show that in the absence of Bez, lipids accumulate in the fat body, the major lipid-storing organ of *Drosophila* with functional homology to liver and adipose tissue.

Lipids are trapped in Bez mutant adipocytes, as becomes apparent by enlarged lipid droplets and by an inability to mobilize lipids in response to starvation. Lipid export from storing tissue is not only important to catabolize lipids for energy gain in times of nutrient scarcity, but also serves to nurse target tissues such as imaginal discs, ovaries and the brain. The requirement of lipids for oocyte development becomes apparent during maturation from stage 8 to stage 10 when oocytes accumulate neutral lipids (Sieber & Spradling, 2015). Bez mutants are viable, but females are sterile and lay very few, collapsed eggs. Bez mutant ovaries degenerate as apparent by the absence of mature eggs and the presence of fragmented DNA in follicles from approximately stage 10 on as well as the missing neutral lipid accumulation. Of note, it is sufficient to deplete Bez from the fat body to reduce fecundity and induce starvation sensitivity. This highlights the importance of lipids stored in adipocytes as dietary lipids directly from the gut cannot substitute. Bez is not expressed in ovaries, which precludes a direct role for Bez in ovary lipidation, but further underlines its role in remote lipid dispatch from the fat body.

Lipid transport through hemolymph is mainly mediated by Lipophorin (Lpp), a lipoprotein with ApoB homology. ApoB is the characteristic apolipoprotein in human chylomicrons and low density lipoprotein (LDL, Olofsson & Borèn, 2005). CD36 can interact with lipoproteins: an important example is the role of CD36 as a receptor for oxidized LDL on macrophages. Upon binding of oxLDL to macrophage CD36 in a blood vessel, the macrophages convert to foam cells and migrate into the intima of the blood vessel, which marks the first step of a cascade leading to the formation of atherosclerotic plaques (Libby, 2021). It is unclear to date if CD36 – lipoprotein interaction also serves nutrient allocation in healthy individuals. We characterize one of the 14 Drosophila SR-Bs as a mediator of storage lipid distribution. To this end, Bez interacts with Lpp. We show that Lpp colocalizes with Bez at the adipocyte plasma membrane. Bez mutant adipocytes can no longer trap Lpp at the plasma membrane. We show in two sets of experiments that lipid transfer from fat body tissue to isolated Lpp is impaired in Bez mutants. BODIPY-stained lipids from wildtype fat body are readily transferred onto exogenous, isolated Lpp; likewise, a traceable oleic acid analog (clickable alkyne oleate) loaded on wildtypic fat body can be detected in Lpp upon incubation with the tissue. We found considerably less BODIPY-stained lipids in Lpp that were incubated with Bez mutant fat body tissue; moreover, while Bez mutant fat body accumulates lipids that incorporated alkyne oleate, they transfer them at a lower rate to Lpp. Of note, we measured this in a time frame of 60 min, which leaves the possibility that lipid transfer is slower, but not completely abolished in Bez mutants.

We propose the following model for the function of Bez: located to the plasma membrane of adipocytes, but also gut cells, Bez mediates the export of lipids. The lipids are transferred to other organs as energy source during nutrient scarcity or energy-demanding developmental processes. For lipid mobilization, Bez interacts with Lpp the major lipid carrier in hemolymph that can also bind to specific receptors in target organs such as oocytes. Lacking Bez, adipocytes fail to suffiently supply lipids. Surprisingly, none of the other 8 SR-B (out of 14 family members) expressed in the fat body (Herboso et al., 2011) can substitute; neither can intestinal Bez. To our knowledge, this study describes a role in lipid export for one of these SR-B for the first time. Other SR-Bs are involved in the uptake of their substrates, e.g. NinaD and Santa-maria in carotenoid uptake or Emp in endocytosis-dependent protein uptake (Pinheiro et al., 2023). Our study suggests that Bez is also required for Lpp interaction and lipid transfer from the gut, since Lpp accumulate in gastric caeca of Bez mutants. While lipid export from gut cells might also be mediated by Bez, it does not affect oocyte maturation. Midgut-specific depletion of Bez neither induces a developmental delay nor ovary degeneration. This puzzling finding may relate to specific characteristics of the mobilized lipids. Moreover, we show a reduced content of medium-chain DG in Lpp, which have been reported to derive from the gut (Palm et al., 2012). The lipid export from gut cells might also be mediated by Bez, but does not affect oocyte maturation. This is demonstrated by midgut-specific depletion of Bez, which neither induces a developmental delay nor ovary degeneration. Dietary lipids are thus insufficient to promote oocyte maturation. We speculate that lipids traffic sequentially and in a coordinated way from the gut to the fat body, and upon remobilization further to different target organs.

Our study opens the paths to several exciting research questions of inter-organ communication: how are lipids sorted and allocated between gut, fat body and target tissues? Tissues like brain and ovaries have completely different lipid compositions; how much does uptake of dietary lipids and remodeling in the fat body contribute to their lipid signature? Is there a buffering system for the incorporation of lipids of a specific chain lengths? For example, will dietary medium- or long chain lipids affect the composition of ovary lipids, or be adjusted in the fat body? The combination of *Drosophila* as a tractable genetic model that allows straightforward *in vivo* and *ex vivo* manipulation of specific tissues with alkyne lipid tracing by high resolution and high content analysis is a promising approach to answer these questions.

## Material and Methods

### Fly stocks, mutant generation and husbandry

The fly stocks used in this study were obtained from the Bloomington *Drosophila* Stock Center (BDSC) at Indiana University: *bez^EP^* (#28496), *w* (#6329), *c355-Gal4* (#3750), *cg-Gal4* (#7011), *lsp2-Gal4* (#6357), *V32-Gal* (#4937), *prd-Gal4* (#1947), UAS-Dcr-2 (#24651), UAS-Flp (#4540), and Ga4.Act5C (FRT.CD2) (#4780), and Gal4-Act5C (FRT.CD2), UAS-RFP (#30558); from the Vienna Drosophila Resource Center (VDRC): the bez-RNAi lines *P(GD2224)v42872* (#42872) and *P(KK111090)VIE-260B* (#103492) and the apoLpp-GFP line *PBac(fTRG00900.sfGFP-TVPTBF)VK00033* (#318255) that expresses apoLpp tagged with sfGFP at the C-terminus; from the National Institute of Genetics (NIG-FLY) at Japan the bez-RNAi line # 3829R-2. The group of Ronald Kühnlein (University of Graz, Austria) kindly provided the stocks: *y*w*; P{w[+mW.hs]=GawB} FB P{w[+m*]UAS- GFP1010T2}#2; P{w[+mC]=tubPGal80[ts]2*; Sb1 e1 and *w*; P{w[*mW.hs]=GawB} FB; +*; and UAS-Plin1-EGFP (w*;P{w+mC UAS-plin1::EGFP} / TM3).

The bez^EP^ allele has the EP-element P{w[+mC]=EP}CG3829[G8378] of 7.987 kb inserted in the third exon of *bez* gene locus, 317 bp downstream of the translation initiation site ATG (Figure 2A). We generate the null bez^jo2^ allele by mobilization of the EP-element that carries a mini-white gene [w^+^]. Candidates were selected by loss of the mini-white transgene and by the sterility phenotype found in homozygous bez^EP^ mutant females. PCR of genomic DNA of the bez^jo2^ allele showed that 35 nucleotides of the EP-element remained inserted in the Bez coding region and introduced a premature termination code after amino acid 88. The apoLpp-GFP line was combined with the *bez^jo2^* allele to yield bez-/-; Lpp-GFP.

The genotype of the bez-RNAi stock used in this study is *w; UAS-bez^3829R-2^; UAS-Dcr2, UAS-bez^GD2224^.* This line was generating by combination of the UAS-Dcr and two the bez RNAi lines: # 3829R-2 from NIG and the bez RNAi line # 42872 from VDRC. Similar, but weaker phenotypes were obtained with both independent RNAi lines, as well as with the VDRC line #103492.

The generation of the conditional bez-RNAi knockdown was done using the TARGET system (McGuire, Mao, & Davis, 2004). The crosses between the Gal4 driver and the bez RNAi flies were done at the restrictive temperature of 18°C, at which the Gal80 protein is active, represses Gal4 and the UAS-bez-RNAi constructs are not expressed. Adult flies were collected in the first 24 hours after eclosion, separated by sex and shifted to 29°C, temperature at which the Gal80 protein is inactive and the UAS-bez-RNAi constructs start to be expressed. The flies were kept at 29°C for 4 days before experiments were performed.

Clones in the fat body were generated by combining prd-Gal4 and UAS-Flp with an Actin-Gal4 STOP cassette (Gal4-Act5C (FRT.CD2)) and UAS constructs. The prd enhancer is active during embryonic development and drives the expression of the flipase that through the excision of a STOP cassette allows the expression of the UAS constructs (the fluorescent reporters UAS-GFP or UAS-RFP and UAS-bez-RNAi for knockdown or UAS-Bez for rescue experiments) independent from the original prd enhancer. The prd-Gal4 has a transient, but very broad, expression pattern in cells that will differentiate into fat body cells, which allow recovering many clones in adipocytes that have been induced very early in development.

### Generation of anti-Bez antibodies and Immunostainings

Anti-Bez antibody was generated after guinea pig immunization with the peptide SHTKDAEMSMPARQESDR-Cys (amino acids: 2-19) at Pineda Antibody service (Berlin). The affinity purified anti-Bez antibody was used at 1:100 dilution. The following additional primary antibodies were used: rabbit anti-GFP (1:100; Santa Cruz), mouse anti-RFP (1:200; Santa Cruz), goat anti-E-cadherin (1:100; Santa Cruz), rabbit anti-plin1 (1:100, gift from R. Kühnlein), mouse anti-α-spectrin (1:10; 3A9, Developmental Studies Hybridoma Bank) and mouse anti-orb (1:100; 4H8 and 6H4 (1:1), Developmental Studies Hybridoma Bank). Conjugated secondary antibodies Alexa488 (Molecular Probes) and Cy3 (Jackson ImmunoResearch) were used at 1:200. Conjugated Alexa647 (Molecular Probes) were used at 1:100. Stainings with DAPI, Nile Red, and BODIPY 558/588_C12_ (Sigma Aldrich, D-3835) were performed for 2 hours at room temperature during the incubation with the secondary antibodies. Embryos were mounted in Fluoromount medium (Southern Biotech). Fluorescent images were obtained on a Zeiss (LSM710) confocal microscope and Airyscan super-resolution were obtained on a Zeiss LSM 880. Quantification of lipid droplet size and fluorescence intensities were done with ImageJ. For colocalization analysis, we used the Plugin JACoP. For quantification of lipid droplet size, we followed the protocol by (Ugrankar et al., 2019). Cells from 5 images from at least 3 different experiments were analyzed.

### Molecular Biology and constructs

For pUAST-Bez generation, Bez coding sequence was amplified using the primers CCGCGGCCGCTGATGTCACATACCAAAGATGCAGAG and GCTCTAGATTACGTTCCCTGATGGATGCCCCCA from the GH19047 cDNA (Drosophila Genomics Resource Center (DGRC)) and inserted in pUAST by using NotI and XbaI. Transgenic flies were generated using standard P-element transformation.

Quantitative real-time RT-PCR was performed with the iQ5 Real-Time PCR Detection System from BIO-RAD, using the IQ SYBR Green Supermix (Biorad). The RNA was extracted using the NucleoSpin RNA II kit (Machery & Nagel) and the cDNA were synthesized using the QuantiTect Reverse Transcription Kit (Qiagen). cDNA samples were run in triplicates and experiments were repeated with independently isolated RNA samples. The mRNA amounts were normalized to *rpL23 (rp49)* mRNA values.

### Quantification of egg deposition

For the quantification of egg depositions, crosses of a UAS-bez*-*RNAi line and different Gal4-driver were performed with 10 females and transferred to 29 °C after 2 days. Eggs were collected in apple juice agar plates coated with a thin layer of yeast in intervals of 24 hours. To determine production rates the total number of eggs was counted over the following two days.

### Starvation assay

For the starvation assay 20 four-day-old flies that were previously separated by sex were transferred to starvation vials containing starvation medium (1 % agar in PBS) that only provide water supply. Mortality rates were scored at 12-hour intervals by counting the number of dead flies as diagnosed by the lack of sit-up response and finally determined as a percent of the total population. The starvation assays with *bez*^EP^ mutants were performed at 25°C and for assays with conditional bez-RNAi flies were maintained at 29°C.

### Triglyceride measurements

Eight to ten adult flies were collected in 2 ml screw caps and frozen at -80°C. They were homogenized in 600 μl PBT using the Precellys 24. After homogenization the tubes were centrifuged at maximal speed at 4°C and the supernatant was transferred to a cold eppendorf tube (if necessary, they were centrifuged a second time to remove debris). 10 μl were removed from the supernatant and stored at -80°C in Eppendorf tubes for protein determination using the Pierce BCA Protein Assay Kit. Protein content was calculated based on the albumin standard curve.

The remaining supernatant was heated for 10 min at 70°C and 20 μl of the heated supernatant, as well as 20 μl glycerol standards (0.125, 0.25, 0.5 and 1 mg/ml) and a PBST blank, were transferred to an eppendorf tubes in duplicates. To measure free glycerol, one of the samples was treated with 20 μl of PBT only and the other sample was treated with 20 μl of a triglyceride reagent (TGR). The TGR contains a lipase, that digest the TAGs and relieves the glycerol backbone. In the next step, all the probes were incubated for 45 min at 37°C. After incubation, the tubes were centrifuged for 3 min at full speed (RT) and 30 μl of the probes and standards were transferred to a clear-bottom 96-well plate and 100 μl of free glycerol reagent were added to each well using a multichannel pipette. The plate was incubated at 37°C for 5 min sealed with parafilm. Air bubbles and condensate were removed by centrifuging the plate. Absorbance at 540 nm was measured using a TECAN plate reader.

To minimize errors, all the samples and standards were performed in triplicates. The average of the triplicates was subtracted by the average of the blank. Triolein-equivalent standard curves were constructed for the TGR-treated and untreated standards. Free glycerol content of the samples was calculated based on these standard curves. Glycerol concentration of the TGR treated samples was subtracted by the concentration of the untreated samples. The difference in glycerol concentration hereby represents the amounts of TAGs that were digested by the enzymatic reaction. Finally, the values were normalized to protein levels.

### Hemolymph/Lipophorin purification and lipid extraction for lipidomics analysis

The hemolymph purification protocol was adapted from (Fernando-Warnakulasuriya & Wells, 1988). 40 third instar larvae were washed extensively in PBS to eliminate the food attached to the cuticle. The larvae were homogenized in 250 µl of cold PBS containing protein inhibitors. The homogenate was kept on ice for 5 min and then centrifuged at 3000 rpm at 4°C during 20 min. The supernatant was transferred to a new tube avoiding taking the floating layer of fat and the cellular debris from the pellet. The supernatant was centrifuged again at 1000 rpm for 20 min at 4°C. The supernatant was transferred to a new tube and incubated immediately with dissected fat body. For lipidomics analysis, lipids of 50 µl freshly isolated Lpp were extracted by addition of 100 µl LC-MS grade water. 500 µl extraction mix (5/1 MeOH/CHCl_3_ (LC-MS grade) + 20 µL OMICS internal standard were added as described (Thiele et al., 2019). Samples were sonicated for 30 min in a bath sonicator and centrifuged for 2 min at 20000 g. The supernatant was decanted into a fresh tube. 300 µl CHCl_3_ and 600 µl 1% acetic acid in LC-MS grade water were added. Samples were shaken manually for 10 s and centrifuged at 20000 g for 5 min. The lower phase was transferred into a fresh tube and dried in a speed-vac (Eppendorf concentrator plus) for 15 min at 45°C and redissolved in 500 µl spray buffer (8/5/1 isopropanol/MeOH/H_2_O (all LC-MS grade) + 10 mM ammonium acetate + 0.1% acetic acid (LC-MS grade)) until mass spectrometry analysis.

### BODIPY transfer to Lipophorin

Fat body tissue from control and Bez mutant larvae was dissected in triplicates in hemolymph-like buffer (HL3A; 115 mM sucrose, 70 mM NaCl, 20 mM MgCl_2_, 10 mM NaHCO_3_, 5 mM KCl, 5 mM HEPES, 5 mM trehalose, pH 7.2 (Paradis et al., 2022)). Tissue was stained with BODIPY for 60 min. Fat body tissue was washed extensively with HL3A to remove residual BODIPY. Lpp were isolated as described in the preceeding paragraph and added 1/10 to the stained fat body. Lpp were incubated for 60 min with the fat body on a nutator and fat body tissue was manually removed. Lpp were enriched by centrifugation at 3000 g for 15 min, mounted in Fluoromount on a microscope slide and analyzed immediately with a Zeiss LSM 710.

### Alkyne lipid transfer analysis by mass spectrometry

A published protocol (Thiele et al., 2012) to preload Drosophila fat body tissue with alkyne oleate was optimized. Click reaction and mass spectrometry analysis were as described (Thiele et al., 2019; Wunderling et al., 2023). Briefly, fat body tissue from control and Bez mutant larvae was dissected in triplicates in hemolymph-like buffer. Tissue was preloaded with 50 µM alkyne oleate (FA 19:1;Y) in HL3A when incubated on a nutator for 60 min. An aliquot was taken to analyze the fat body lipidome and quantify alkyne oleate uptake and metabolism. The remaining tissue was washed extensively with HL3A to remove adherent alkyne lipid tracer. Isolated Lpp (see above) were added 1/10 to the preloaded fat body tissue and incubated together on a nutator for 60 min. The fat body tissue was carefully removed, first manually and then by passaging through a cell strainer. Flow-through was centrifuged at 4°C and 3000 g for 15 min and the supernatant was removed. Lipids were extracted as described above but additional internal standards for alkyne labelled metabolites were included (Thiele et al., 2019). After drying of the sample, lipids were resolved in 10 µl CHCl_3_ and sonicated for 5 min. 40 µl of C171 click mix (prepared by mixing 10 µl of 100 mM C171 in 50% MeOH (stored as aliquots at −80°C) with 200 µl 5 mM Cu(I)AcCN_4_BF_4_ in AcCN and 800 µl ethanol) were added and the samples were incubated at 40°C overnight (16 h). 200 µl CHCl_3_and 200 µl water were added and the samples were centrifuged at 20000 g for 5 min. The upper phase was discarded and samples were dried in a speed-vac for 15 min at 45°C. Samples were resolved in 500 µl spray buffer and analyzed by mass spectrometry.

### Statistics

Bar charts represent mean and standard deviation. Error bars in curves represent standard error of the mean (SEM). Boxes in box plots represent the interquartile range and median, whiskers represent minimum and maximum. Green squares in box plots represent single data points. We used Microsoft Excel and GraphPad Prism for bar charts and curves and Origin Pro 8G for box plots. We used the software GraphPad Instat for our statistical analyses. Two-sided Student’s t-test was applied for normally distributed data in single comparisons, assuming heteroschedasticy. One-way ANOVA with Tukey-Kramer post-test was used for multiple comparisons. The Kolmogorov-Smirnow test was applied to test normality, and Bartlett’s method was used to test for equal standard deviations within groups. Asterisks represent * = p < 0.05, ** = p < 0.01, *** = p < 0.001. A minimum of 3 biological replicates was used for each analysis.

## Author contribution

Conceptualization: PC, LK, MHB; Methodology: PC, JO, LK, MHB; Investigation: PC, JO, YJ, KSR, LK, MHB; Writing – Original Draft: PC, JO, MHB; Writing – Review & Editing: LK, PC, MHB; Visualization: PC, MHB; Funding Acquisition: MHB; Supervision: PC, MHB.

## Acknowledgments

We thank Darla Dancourt Ramos and Valeria Gulayeva for help with experiments and Almut Wingen and Dominic Gosejacob for discussion. We thank Christoph Thiele for providing the alkyne oleate. We thank Ronald Kühnlein for sharing the Plin1-GFP and Fat body-Gal4 fly line and Plin1 antibody.

## Supplement

**Supplemental Figure S1:**
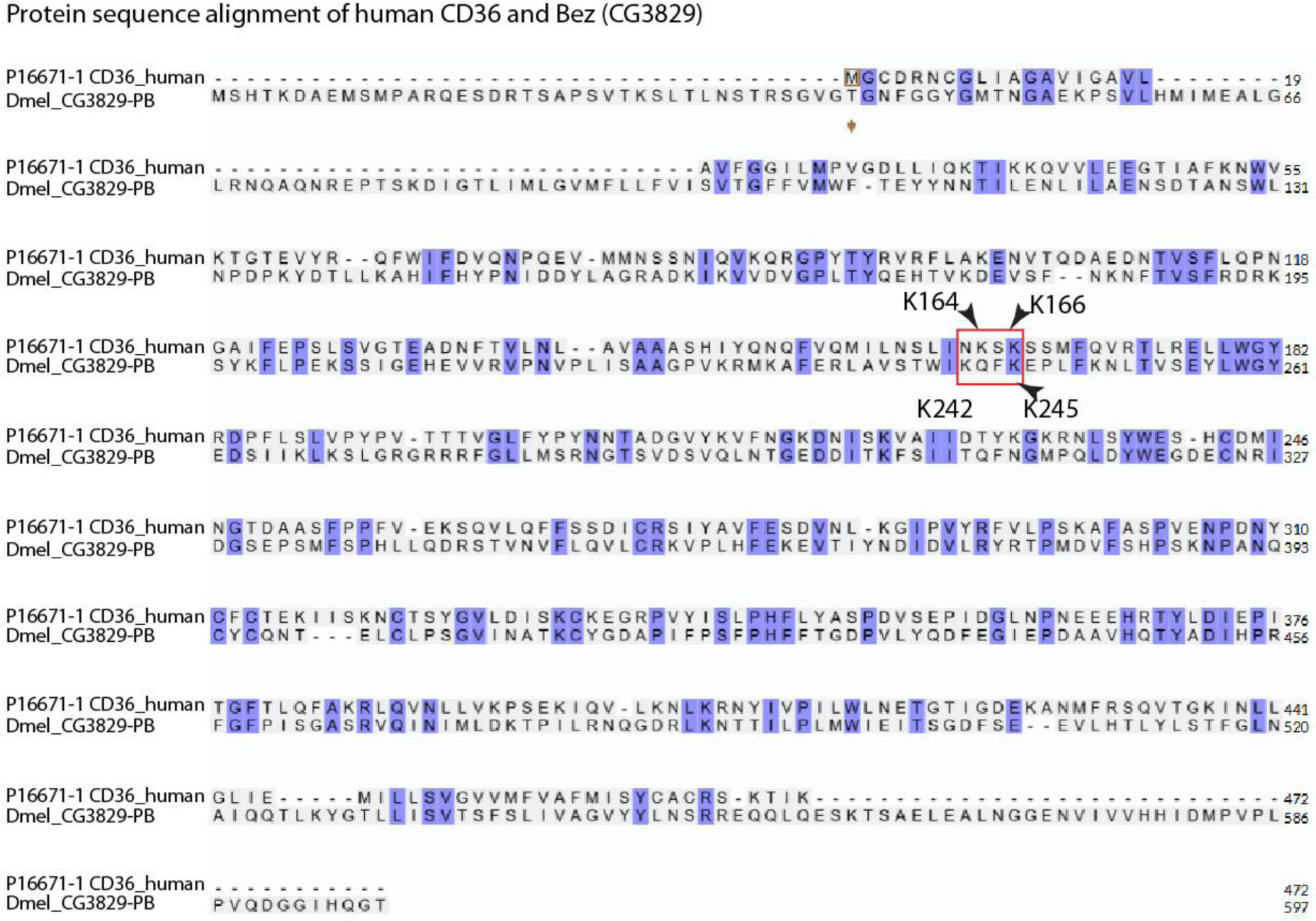
Protein sequence alignment of human CD36 and *D. melanogaster* CG3829 shows several conserved regions and the additional N- and C-terminal amino acid stretches of Bez. Lysine residues of the lipid binding pocket are marked in red.

**Supplemental Figure S2:**
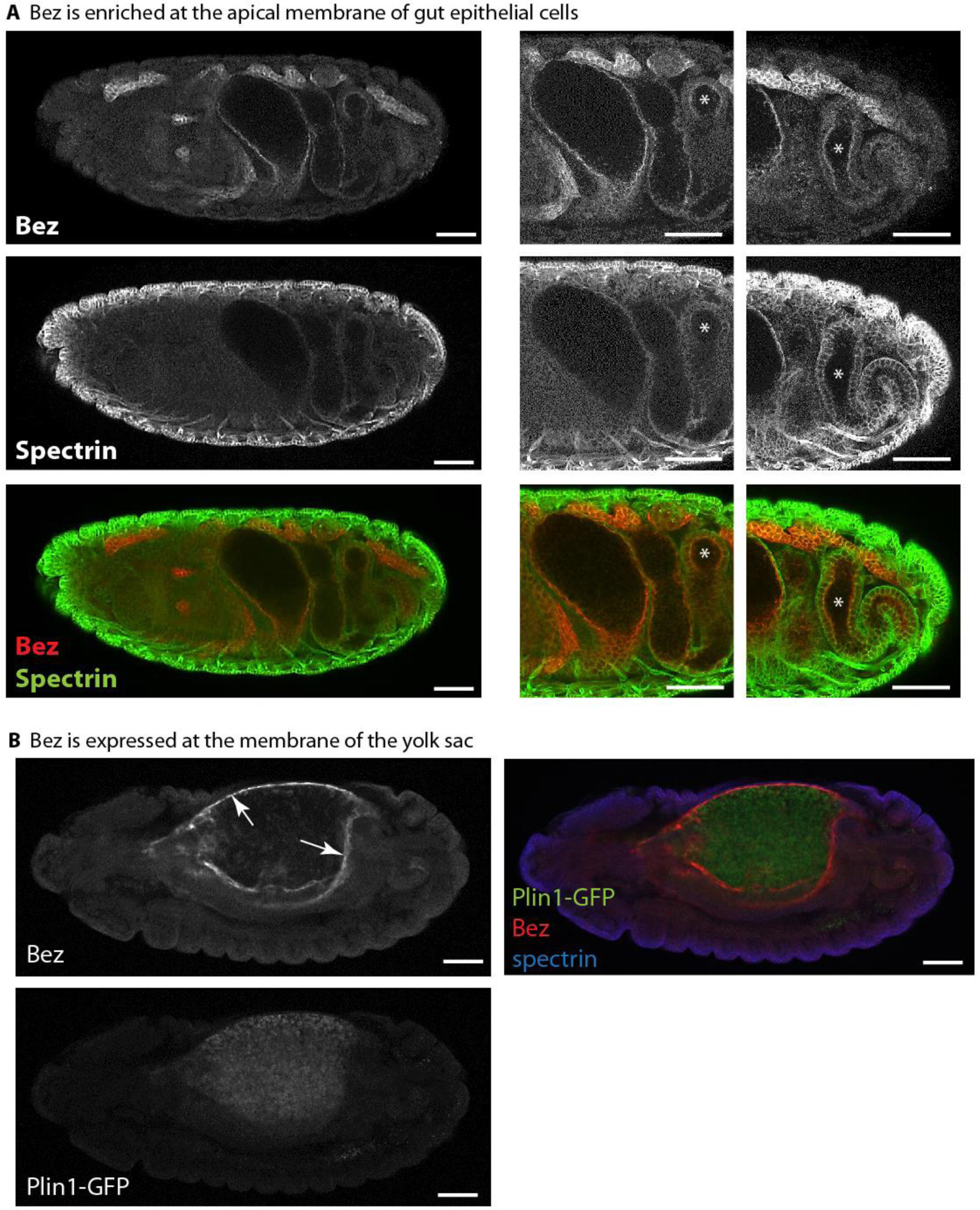
A) In addition to embryonic fat body, Bez is expressed in the gut, where it is enriched in the apical membrane. B) Bez is expressed at the yolk sac. Scale bars represent 50 µm.

**Supplemental Figure S3:**
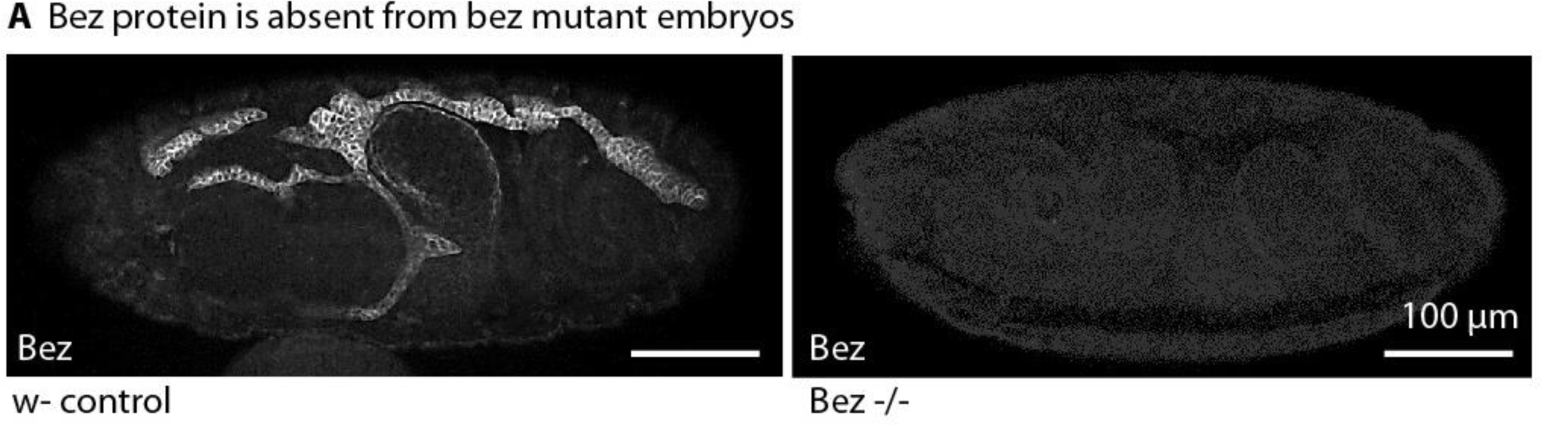
Bez protein is absent from Bez mutant embryos. Confocal images show wildtype and Bez mutant embryos stained with anti-Bez. Scale bars represent 100 µm.

**Supplemental Figure S4:**
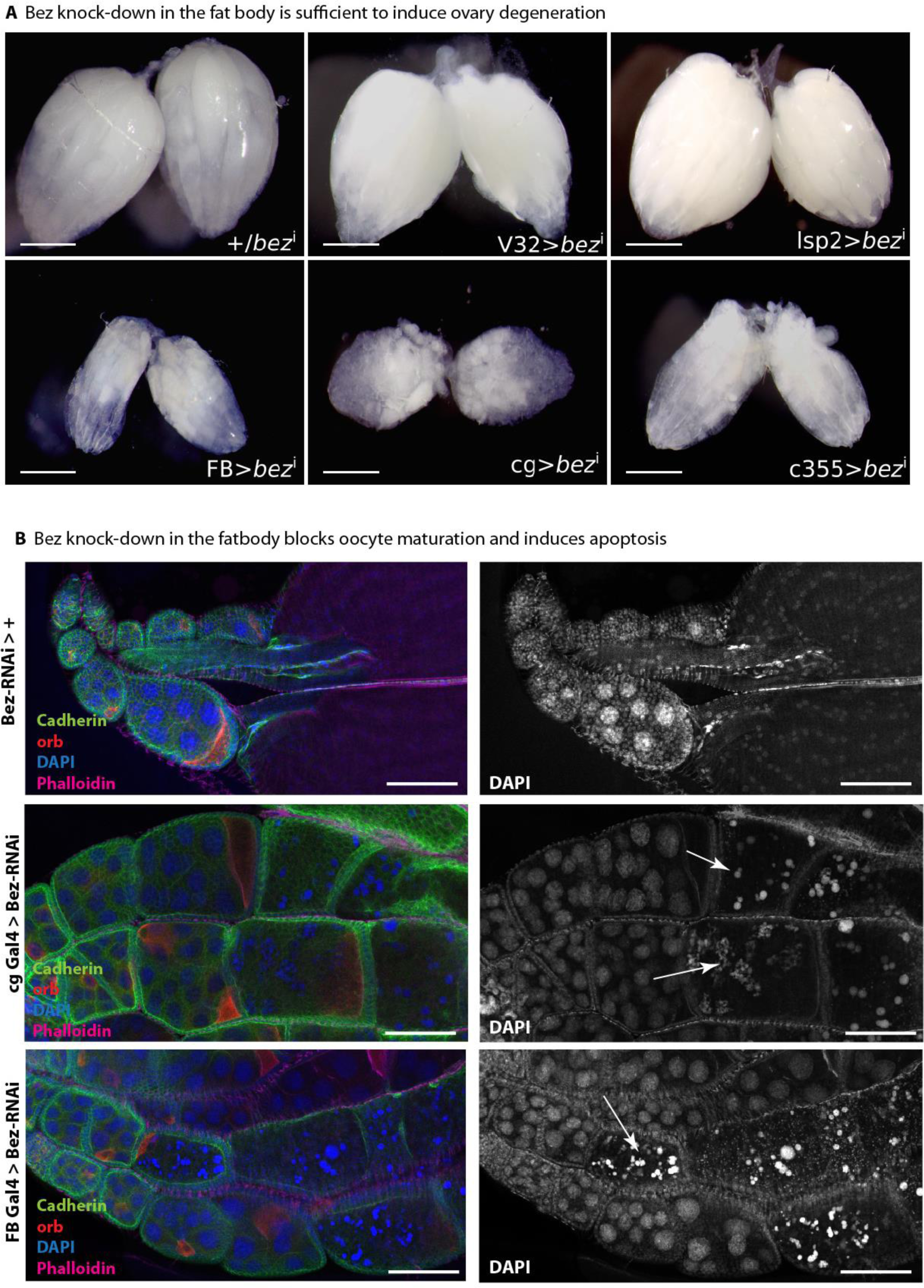
A) bez-RNAi-mediated knock-down is sufficient to induce ovary degeneration. Upper panel from left to right: control without driver; Germline-specific driver (*V32*-Gal4) or hemolymph-specific driver (*lsp2*-Gal4) do not induce ovary degeneration. Lower panel: Fat body-specific knock-down of Bez (*FB*-Gal4, *cg*-Gal4, *c355*-Gal4) induces egg chamber degeneration. B) Knock-down of Bez by RNAi in the fat body (*FB*-Gal4 and *cg*-Gal4) blocks oocyte maturation and induces apoptosis. Mature eggs are absent in fat body-specific bez-RNAi (2^nd^ and 3^rd^ panel) and DNA is fragmented (arrows). Representative images from 5 independent replicates. Scale bars represent 100 µm.

**Supplemental Figure S5:**
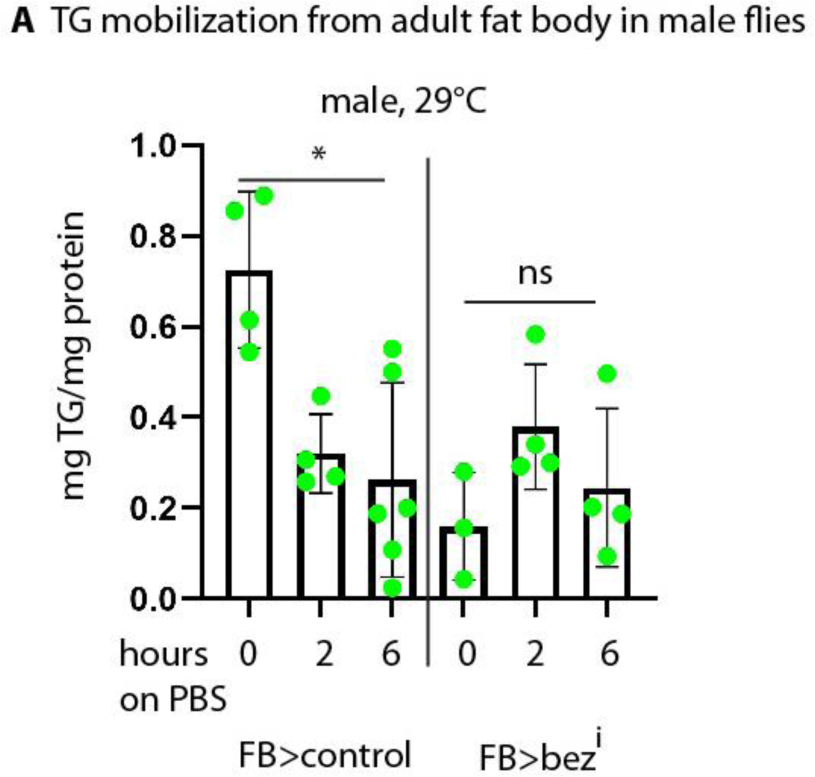
A) Triacylglycerol (TG) as determined by a colorimetric assay from whole male flies expressing bez-RNAi in the fat body (*FB*-Gal4) under the control of the inducible Gal4/Gal80 system. At 29 °C, Gal4 is active and drives the expression of bez-RNAi (right side). In controls, FB Gal4/Gal80^ts^ was crossed to wildtype (left side). Depletion of Bez from the fat body blocks TG reduction upon starvation in male flies. n ≥ 3 in groups of 8 individuals. Asterisks represent * p > 0.05, ns: not significant.

**Supplemental Figure S6:**
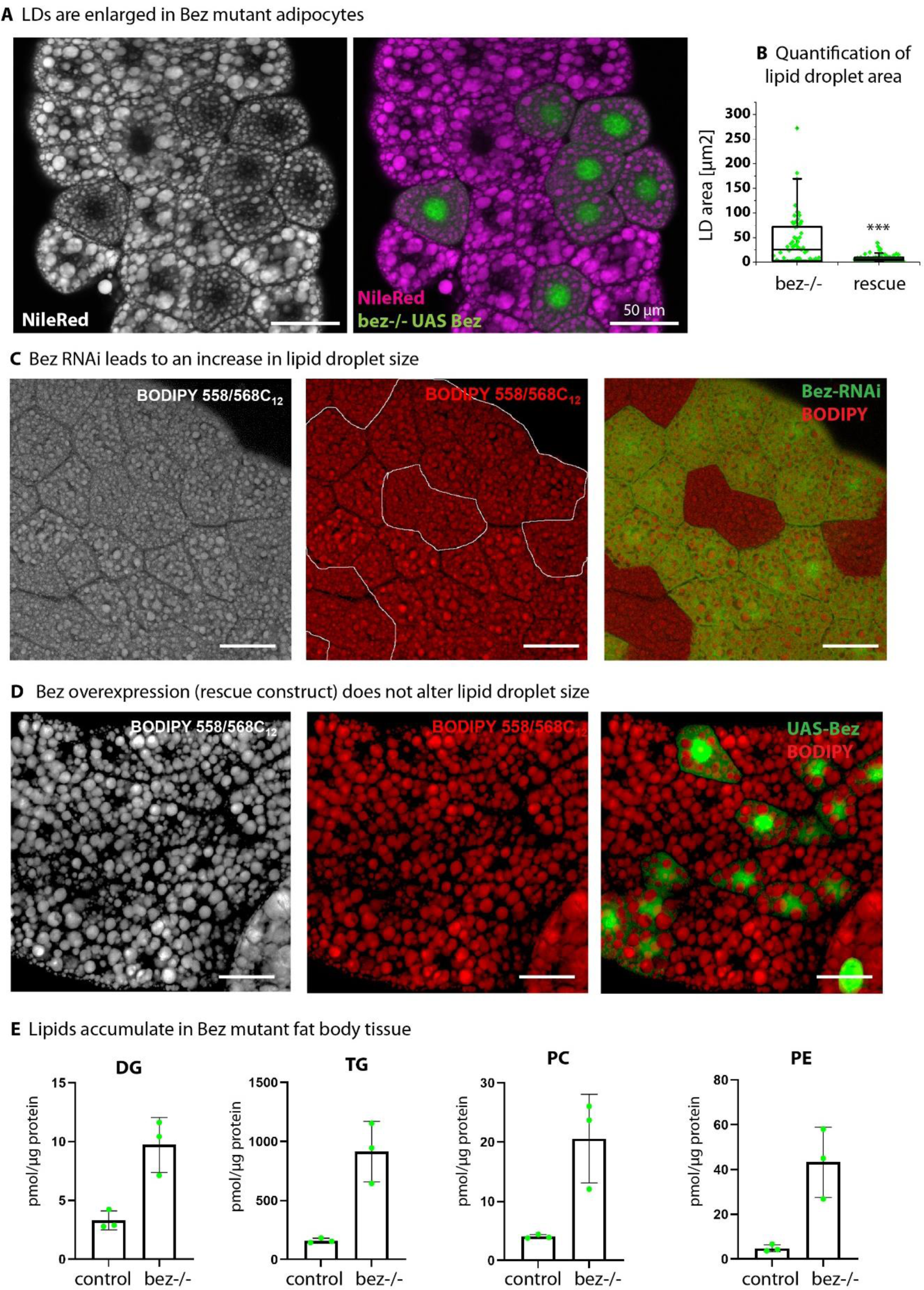
A) Confocal images show Bez mutant adipocytes with rescue clones marked with GFP. Lipid droplets are stained with NileRed. Bez mutant cells show increased lipid droplet size. Representative images from 10 independent replicates. B) Quantification of lipid droplet size (area). Asterisks represent *** p > 0.001. C) Fat body clones expressing Bez RNAi show increased LD size similar to bez-/- adipocytes. Lipids stained with BODIPY. Scale bars represent 50 µm. D) Bez overexpression in a wildtype background has no effect on LD size. Lipids stained with BODIPY. Scale bars represent 50 µm. E) Selected lipid classes as determined by mass spectrometry. DG: diacylglycerol, TG: triacylglycerol, PC: phosphatidylglycerol, PE: phosphatidylethanolamine.

**Supplemental Figure S7:**
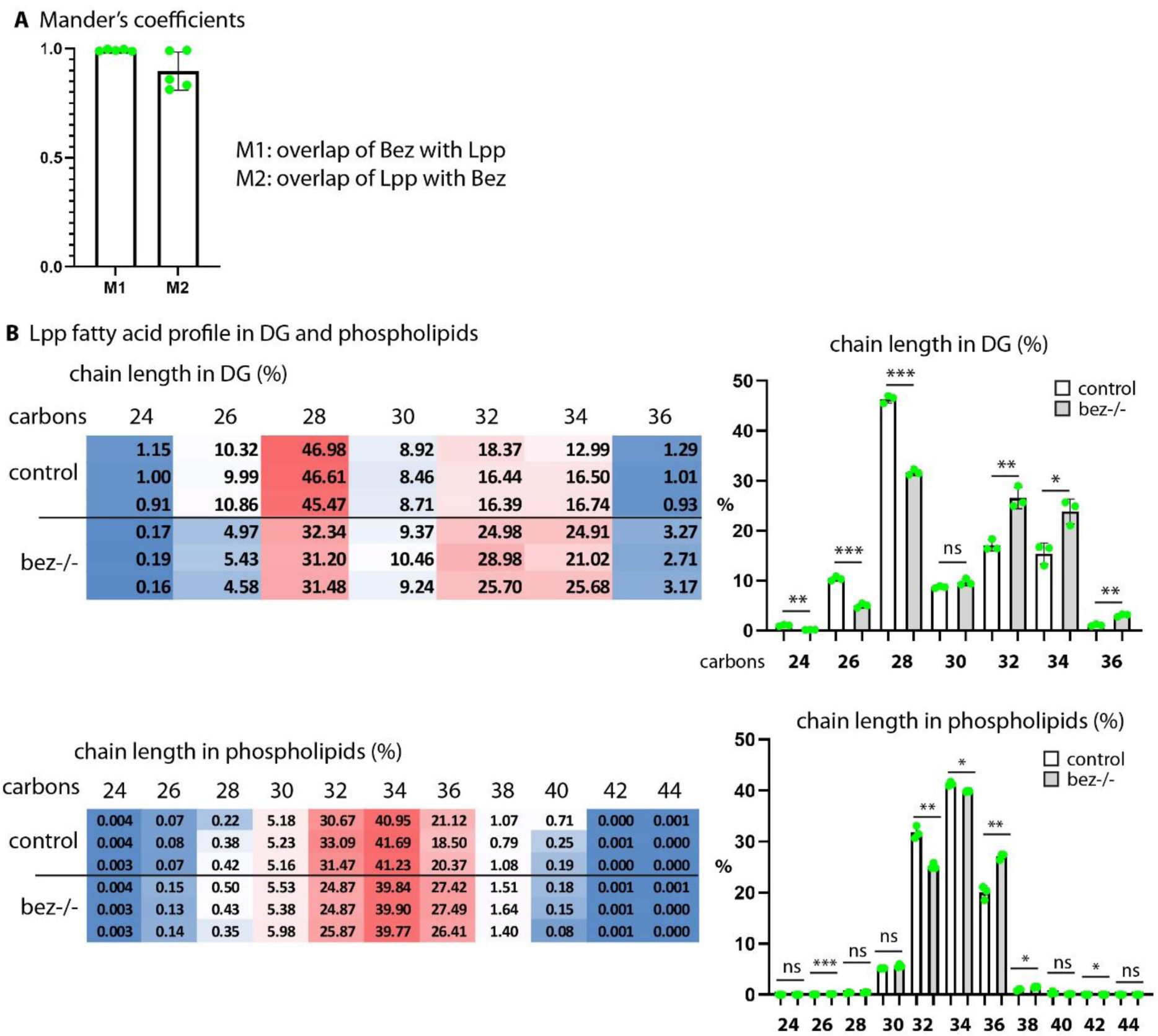
A) Mander’s coefficient for colocalization analysis of Bez with Lpp-GFP (M1) and Lpp-GFP with Bez (M2). B) Fatty acid profile of the Lpp fraction as determined by mass spectrometry. Heat maps show single measurements representing the percentage of species with the indicated chain length. Bar charts show the same data. DG with a combined chain length of 28 carbons (14 carbons average chain length) are predominant in wildtype Lpp, but shifted towards longer chains in Bez mutants. n = 3. Asterisks represent *** p > 0.001, ** p > 0.01, * p > 0.05, ns: not significant.

